# Adaptive evolution of *Pseudomonas putida* in the presence of fluoride exposes moonlighting transporter functions

**DOI:** 10.1101/2025.10.13.682028

**Authors:** Lea Ets, Heili Ilves, Tanel Ilmjärv, Òscar Puiggené, Pablo Iván Nikel, Maia Kivisaar

## Abstract

Fluoride (F^-^), the anionic form of fluorine and the 13th most abundant element in Earth’s crust, is toxic to most organisms above relatively low threshold concentrations. Environmental bacteria often tolerate elevated fluoride levels, but the only known resistance mechanism so far involves CrcB-mediated efflux. In the environmental bacterium *Pseudomonas putida*, CrcB export is the primary defense against fluoride stress. Yet, spontaneous NaF-tolerant mutants emerge even without this transporter, suggesting the existence of additional pathways. To uncover these mechanisms, we performed a genome-wide screen of over 141,000 transposon mutants. We identified PP_3125, a Cro/cI-type transcriptional regulator, as essential for high fluoride tolerance in a Δ*crcB* background. Transcriptomic and proteomic analyses revealed PP_3125-regulated genes, including the benzoate transporter BenE-I, which contributes directly to fluoride tolerance. These findings demonstrate that bacterial transporters can acquire moonlighting functions beyond their canonical roles and reveal previously unrecognized fluoride tolerance strategies in *P. putida*. Together, our results expand understanding of microbial adaptation to toxic ions and provide new targets for engineering stress-resilient strains for environmental and industrial applications.

**IMPORTANCE:** Our work identifies a new fluoride tolerance mechanism in *Pseudomonas putida* that functions independently of the well-characterized CrcB efflux system. We show that inactivation of transcriptional regulator, PP_3125, activates a transporter with an unexpected moonlighting role in fluoride tolerance, highlighting how bacteria can repurpose existing functions to survive environmental stress. This discovery deepens our understanding of microbial stress responses and suggests strategies to engineer robust microbial strains capable of thriving in fluoride-contaminated settings. Such strains could be valuable for bioremediation, sustainable bioprocessing, and other biotechnological applications where fluoride exposure limits microbial performance.

## 1. Introduction

Microorganisms frequently encounter environmental stressors, including heavy metals, organic solvents, and toxic anions, which necessitate various adaptive mechanisms for survival and growth. One key aspect of microbial resilience is chemical tolerance—the capacity of cells to detect, mitigate, and adapt to toxic compounds. Bacteria have evolved diverse strategies to cope with such stresses, including the activation of efflux transporters, modification of membrane permeability, enzymatic detoxification, and reprogramming of cellular metabolism (1,2). Understanding these mechanisms has significant implications for biotechnology, remediation of environmental pollutants, and synthetic biology, where engineered microbes are often exposed to non-natural or stressful substrates.

Fluoride, one of the most common environmental halogens, is a pervasive toxicant to microbial life. At sublethal concentrations, fluoride disrupts metabolic processes by inhibiting key enzymes, such as enolase and pyrophosphatases, often by forming complexes with essential metal cofactors (3,4). To counteract this, many microorganisms possess fluoride-specific ion transporters, the best known of which is the CrcB protein. CrcB functions as a fluoride exporter and is considered a primary defense mechanism against fluoride toxicity in bacteria, including *Escherichia coli*, *Bacillus subtilis*, and *Pseudomonas* species (4–6).

*Pseudomonas putida* is a Gram-negative environmental bacterium renowned for its metabolic versatility, solvent resistance, and capacity to degrade various xenobiotic compounds (7). Its physiological robustness has led to its widespread adoption as a model organism for environmental and industrial biotechnology. *P. putida* KT2440 is a well-characterized, genetically tractable strain chassis for metabolic engineering (8). One emerging area of interest is the use of *P. putida* in biofluorination—the biosynthesis of organofluoride compounds, which are otherwise chemically challenging to produce (6,9,10). This process often involves exposure to sodium fluoride (NaF) as a substrate or intermediate, underscoring the importance of engineering fluoride-tolerant microbial platforms (4,6,11).

Chemical tolerance in bacteria is frequently governed by transcriptional regulators that orchestrate the cellular stress response. Negative regulators can suppress the expression of tolerance-conferring genes, and their inactivation often leads to increased tolerance. One such regulator is PsrA. This transcriptional repressor regulates fatty acid metabolism. In its absence, the growth rate and polyhydroxyalkanoate (PHA) production are reduced, indicating its role in negatively regulating genes related to fatty acid metabolism (12). Additionally, multidrug and metabolite efflux transporters are known to contribute to the export of toxic compounds, including ions and solvents (2,13). Such systems may play a role in fluoride tolerance, although their specific involvement remains understudied. As of yet, CrcB-mediated export is known to be the primary mechanism of fluoride tolerance in *P. putida*.

In this study, we report that evolution of a Δ*crcB* derivative of *P. putida*, lacking the fluoride efflux pump, can give rise to spontaneous NaF-tolerant mutants, suggesting the existence of secondary or compensatory tolerance mechanisms. Through a combination of spontaneous NaF-tolerant mutant selection, transposon mutagenesis, targeted gene deletions, proteomic profiling, and transcriptomic analysis, we identified a previously uncharacterized transcriptional regulator, PP_3125, as a key modulator of fluoride sensitivity. We further demonstrated that inactivation of *PP_3125* derepresses adjacent genes, including the benzoate transporter BenE-I, which contributed to elevated fluoride tolerance.

## 2. Materials and methods

### 2.1 Bacterial strains, plasmids, and culture media

Bacterial strains and plasmids used in this study are listed in Table 1. *P. putida* strains are derivatives of KT2440 (14). Bacteria were grown in lysogeny broth (LB). *E. coli* was incubated at 37°C, and *P. putida* at 30°C. When selection was needed, the growth medium was supplemented with antibiotics at the following concentrations: kanamycin 50 μg · ml^−1^, ampicillin 100 μg · ml^−1^, and gentamycin 10 μg · ml^−1^. Bacteria were electrotransformed according to the protocol of Sharma and Schimke (15). Sodium fluoride (NaF) was purchased from Sigma-Aldrich (St. Louis, MO, USA; cat. # 201154) and used in various concentrations, between 0 and 30 mM, specified in the text.

**Table 1.** Strains and plasmids used in this study.

| Bacterial strain or plasmid | Relevant characteristics | Source or reference |
| --- | --- | --- |
| *Escherichia coli* |  |  |
| DH5α λpir | Cloning host; F<sup>-</sup>λ<sup>-</sup>endA1 glnX44(AS) thiE1 recA1 relA1 spoT1 gyrA96(Nal<sup>R</sup>) rfbC1 deoR nupG Φ80(lacZΔM15) Δ(argF-lac)U169 hsdR17(r<sub>K</sub><sup>-</sup>m<sub>K</sub><sup>+</sup>), λpir lysogen | (21) |
| CC118 λpir | Cloning host; $\Delta(ara-leu)$ *araD* $\Delta lacX174$ *galE* *galk* *phoA* *thiE1* *rpsE* *rpoB*(Rif<sup>R</sup>) *argE*(Am) *recA1*, λpir lysogen | (22) |
| HB101 | Helper strain; F<sup>-</sup> $\lambda$ *hsdS20*(*r<sub>B</sub><sup>-</sup> m<sub>B</sub><sup>-</sup>*) *recA13* *leuB6*(Am) *araC14* $\Delta$(*gpt-proA*)62 *lacY1* *galk2*(Oc) *xyl-5* *mtl-1* *thiE1* *rpsL20*(Sm<sup>R</sup>) *glnX44*(AS) | (23) |
| *Pseudomonas putida* |  |  |
| KT2440 | Wild-type strain; mt-2 derivative cured of the TOL plasmid pWW0 | (24) |
| Δ*crcB* | KT2440 Δ*crcB* | (6) |
| Δ*crcB*Δ3125 | KT2440 Δ*crcB* ΔPP_3125 | This study |
| Δ3125 | KT2440 ΔPP_3125 | This study |
| Δ*crcB*Δ3125_tac_3125 | KT2440 Δ*crcB*Δ3125 strain containing *lacI*-P<sub>tac</sub>-PP_3125 in the intergenic region between *glmS* and PP5408 (Gm<sup>r</sup>) | This study |
| Δ*crcB*Δ5 | KT2440 Δ*crcB* ΔPP_2033, ΔPP_2034, Δ*benE-I*, ΔPP_2036 ΔPP_2037 | This study |
| *crcB1* | Spontaneous NaF<sup>+</sup> Δ*crcB* mutant with genomic deletion [PP_2985]-PP_5598 | This study |
| *crcB4* | Spontaneous NaF<sup>+</sup> Δ*crcB* mutant with genomic deletion [PP_2985]-[PP_3288] | This study |
| *crcB12* | Spontaneous NaF<sup>+</sup> Δ*crcB* mutant with genomic deletion [PP_3119]-[PP_3133] | This study |
| *crcB18* | Spontaneous NaF<sup>+</sup> Δ*crcB* mutant with genomic deletion [PP_3059]-[PP_3221] | This study |
| Plasmids |  |  |
| pBAM1 | Delivery plasmid for mini-Tn5 (Amp<sup>r</sup>, Km<sup>r</sup>) | (25) |
| pRK2013 | Helper plasmid for conjugal transfer (Km<sup>r</sup>) | (26) |
| pSEVA/lacI-tac | Broad-host range plasmid containing *lacI*-P<sub>tac</sub> cassette (Km<sup>r</sup>) | (27) |
| pSNW2 | pEMG derivative with P14g(BCD2)→msfGFP (Km<sup>r</sup>) | (17) |
| pSW(I-SceI) | Plasmid for I-SceI expression (Amp<sup>r</sup>) | (16) |
| pSNW2_3125_before | pSNW2 plasmid containing a fragment of 500 nucleotides before the PP_3125 gene with BamHI and SacI restriction sites (Km<sup>r</sup>) | This study |
| pSNW2_3125_before+after | pSNW2 plasmid containing fragments of 500 nucleotides before and 500 nucleotides after the PP_3125 gene, with a SacI restriction site between the two fragments (Km<sup>r</sup>) | This study |
| pSEVA/lacI-tac-3125 | pSEVA/lacI-tac containing gene cassette *lacI*-P<sub>tac</sub>-PP_3125 (Km<sup>r</sup>) | This study |
| pGP704LTn7GmI-lacI-tac-3125 | pGP-miniTn7-ΩGm containing gene cassette *lacI*-P<sub>tac</sub>-PP_3125 between inverted repeats of mini-Tn7 (Amp<sup>r</sup>, Gm<sup>r</sup>) | This study |
| pSNW2/del5 | pSNW2 plasmid containing fragments of 500 nucleotides after PP_2033 and 500 nucleotides after PP_2037 gene (Km<sup>r</sup>) | This study |
| pSEVA/lacI-tac-*benE-I* | pSEVA/lacI-tac containing gene cassette *lacI*-P<sub>tac</sub>-*benE-I* | This study |
|  | (Km<sup>r</sup>) |  |
| pSEVA/lacIac-2034 | pSEVA/lacIac containing gene cassette *lacI*-P<sub>tac</sub>-*PP_2034* (Km<sup>r</sup>) | This study |
| pGP-miniTn7-ΩGm | pGP704 L carrying a SacI-XbaI mini-Tn7-ΩGm cassette from pBK-miniTn7-ΩGm (Amp<sup>r</sup>, Gm<sup>r</sup>) | (18) |
| pUX-BF13 | Helper plasmid, providing the Tn7 transposase proteins (Amp<sup>r</sup> *mob*<sup>+</sup>) | (20) |

### 2.2 Construction of plasmids and strains

Oligonucleotides used in this study are listed in Table 2. For the construction of *the PP_3125* deficient strain, 500 nucleotides of the upstream and downstream region of *PP_3125* were amplified from chromosomal DNA of *P. putida* KT2440 with oligonucleotide*s* PP_3125_BHI and PP_3125delSac for the upstream region and PP_3125outdelSac and *PP_3125*_EcoRI for the downstream region. The PCR product of the upstream region of PP_3125 was cloned into BamHI- and SacI-opened pSNW2, resulting in pSNW2_*3125*_before. Next, the pSNW2_*3125*_before was opened with SacI and EcoRI, and the downstream region of *PP_3125* was cloned into the vector, resulting in the vector pSNW2_*3125*_before+after. A combination of previously published methods was used to carry out the gene deletion on *P. putida* strains (16,17). The plasmid pSNW2_*3125*_before+after was electroporated into *P. putida* KT2440 WT and Δ*crcB* strains. Kanamycin-resistant colonies with visible GFP expression under blue light, carrying a cointegrate in the chromosome, were isolated on kanamycin selective plates and then were electroporated with the I-SceI expression plasmid pSW(I-SceI). To resolve cointegrates, the plasmid-encoded I-SceI nuclease was induced by cultivating bacteria overnight in LB medium supplemented with 1.5 mM 3-methylbenzoate. Kanamycin-sensitive colonies were isolated, and PCR and DNA sequencing verified the PP_3125 deletion. The deletion strains were cured of the plasmid pSW(I-SceI) by overnight growth without antibiotics. The absence of pSW(I-SceI) was confirmed by PCR.

**Table 2.**
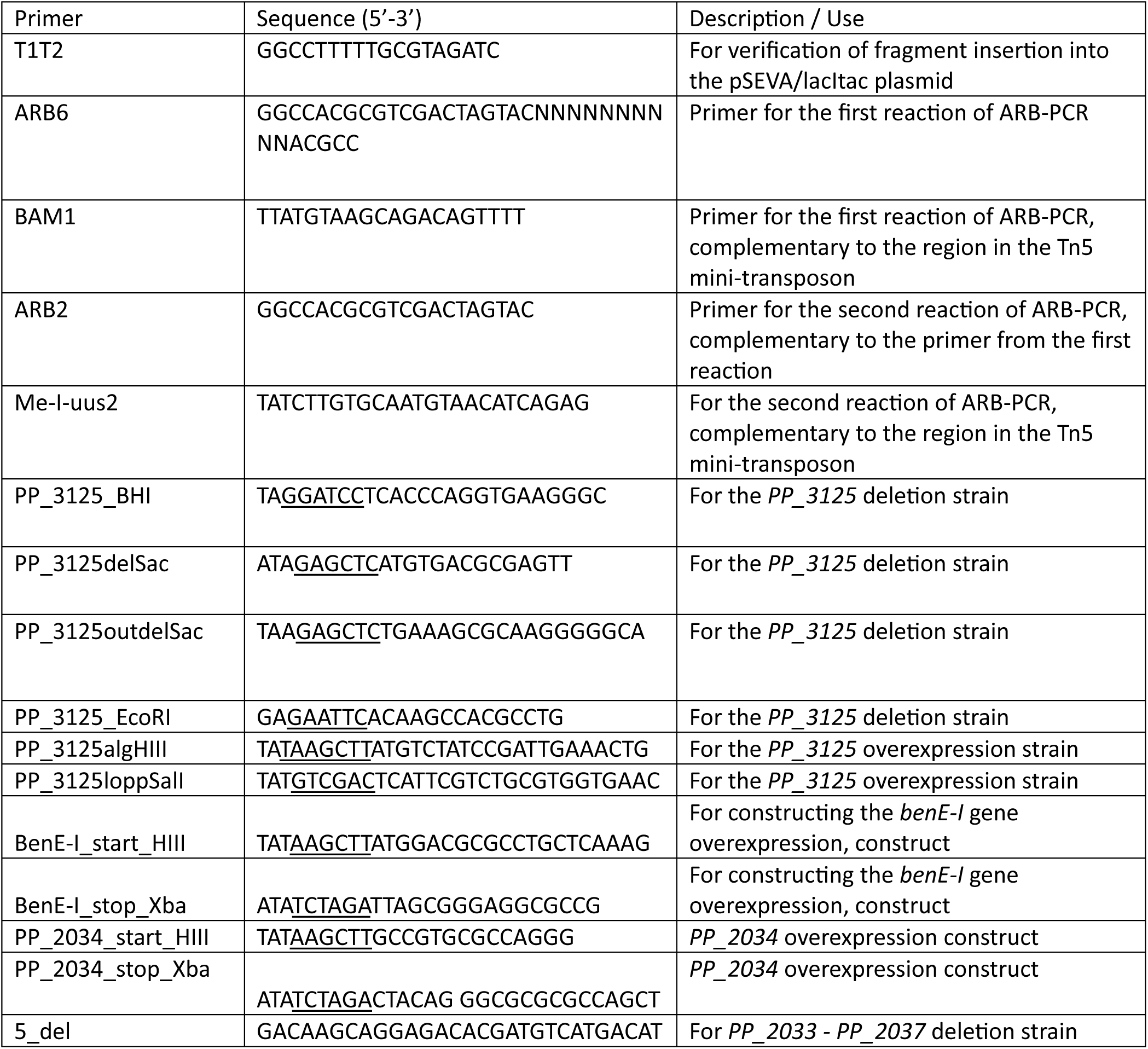
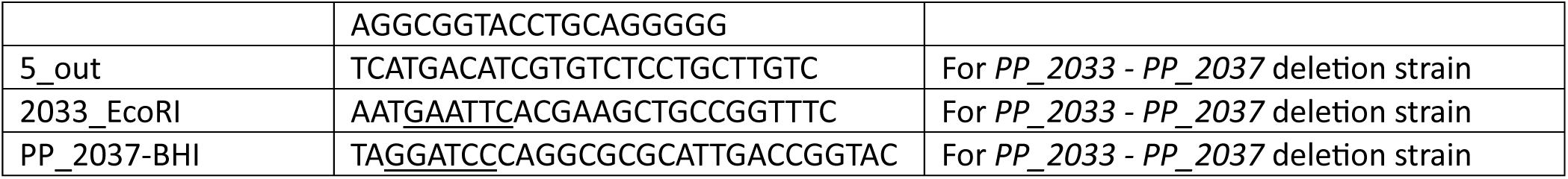
Primers used in this study.

For the construction of the *PP_2033-PP_2037* operon-deficient strain, a combination of previously published methods was used (16,17). Approximately 500 bp from the upstream and downstream regions of the operon were separately amplified and then fused into one PCR fragment by overlap extension using the primer pair 2033_EcoRI and PP_2037-BHI. The obtained PCR fragment (∼1 kb) was cleaved with EcoRI and BamHI and cloned into the corresponding sites of the plasmid pSNW2. The following steps were the same as for the *PP_3125* deletion strains described above.

For the construction of overexpression constructs, the genes *PP_3125, PP_2034, and benE-I* were amplified by PCR from the chromosome and inserted into the vector plasmid pSEVA/lacItac by using the HindIII and SalI restriction sites for *PP_3125*, and HindIII and XbaI for *PP_2034* and *benE-I,* resulting in pSEVA/lacItac-*3125,* pSEVA/lacItac-*benE-I,* or pSEVA/lacItac-*2034*, respectively. To construct the overexpression strain of *PP_3215,* the whole gene cassette from pSEVA/lacItac-*3125* was cloned into pGP-miniTn7-ΩGm (18) vector as a NotI fragment, resulting in plasmid pGP704LTn7GmlacItac-*3125*. All previous steps were carried out in *E. coli*. For the delivery of the gene cassette into the Tn7 insertion site in the chromosome of *P. putida,* the previously published method was used (19). *P. putida* Δ*crcB*Δ*3125* strain was co-electroporated with plasmid pGP704LTn7GmlacItac-*3125* and helper plasmid pUX-BF13 (20). The positive clones on LB Gm plates were identified by PCR and verified by sequencing.

### 2.3 Estimation of the frequency of spontaneous NaF-tolerant mutants in the ***Δ****crcB* strain

To analyze the occurrence of spontaneous NaF-tolerant mutants in the Δ*crcB* strain, bacteria were grown for 6 hours in LB liquid medium, a 10^-5^ dilution of the cultures was made, dispensed into at least 10 test tubes as 2.3 ml aliquots, and allowed to reach saturation by growing cells for 18 h. ∼3×10^9^ cells were plated onto LB plates containing 5 mM NaF and incubated at 30 °C. The NaF^+^ colonies were counted on day 5. The frequency of NaF^+^ colonies is the number of NaF^+^ colonies per milliliter divided by CFU. Four biological replicates with at least 10 parallels were analyzed.

### 2.4 Whole genome sequencing

For whole genome sequencing, DNA from spontaneous NaF+ Δ*crcB* mutants was extracted using the Thermo Scientific GeneJET Genomic DNA Purification Kit. Sequencing was performed on the Illumina MiSeq platform with 100x coverage in the University of Tartu. All the resulting raw sequences are available in GenBank under BioProject ID PRJNA1330567. Received raw data was trimmed using Trimmomatic ver. 0.39 (28). Mutations in the sequences were detected with Breseq ver. 0.36.0 (29). Changes in the chromosome are mapped against the published genome of *Pseudomonas putida* KT2440 (7).

### 2.5 Transposon mutagenesis and selection of transposon mutants

A transposon mutagenesis of *P. putida* Δ*crcB* was performed to identify the genes affecting fluoride tolerance. Mini-transposon Tn5 (Km^r^)-carrying plasmid pBAM1 (25) replicating only in *E. coli* hosts containing the phage lambda *pir* gene (e.g., *E. coli* DH5α λ*pir*) was transferred into the Δ*crcB* by conjugation with the aid of the helper plasmid pRK2013 (26). Transconjugants of Δ*crcB* carrying random insertions of mini-Tn5 in the chromosome were selected on LB-Km plates. The total number of transposon mutant colonies was counted, and the colonies were transferred to LB plates containing 15 mM NaF using replica plating. About 141 000 transposon mutants from three independent transposon mutagenesis experiments were obtained, out of which 48 had a fluoride-tolerant phenotype. The NaF-tolerant colonies were isolated and subjected to the determination of the location of the mini-Tn5 insertion in the chromosome by arbitrary PCR and DNA sequencing.

### 2.6 Arbitrary PCR

To identify the mini-Tn5 insertion site in the chromosome of *P. putida* Δ*crcB* transposon mutants, an arbitrary PCR and DNA sequencing were performed. The arbitrary PCR products were obtained according to the published protocol (30) by two rounds of amplification. In the first round of PCR, primers BAM1 and arbitrary primer ARB6 were used. The second-round PCR product was generated with primers ME-I-uus2 and arbitrary primer ARB2. DNA sequencing of the PCR products was performed using the BigDye Terminator v3.1 Cycle Terminator kit and analyzed with the Applied Biosystems 3730×l DNA sequencer.

### 2.7 Growth analysis experiments

For growth analysis experiments, the cells were grown overnight in liquid LB medium, then diluted to an OD600 of 0,05 and NaF was added at a final concentration of 0 mM up to 2,5 mM before transferring 100 µL of the diluted culture to the 96-well plate. The optical density was measured automatically every 10 minutes for 24 hours. Data were analyzed, and graphs and growth measurements were obtained from a QurvE program version. 1.1 (31). Statistical analysis was done by using GraphPad Prism 10.0. D′Agostino & Pearson test was used to determine the normality of the dataset. As the data did not follow a normal distribution, the non-parametric method was used to compare different datasets. A Kruskal-Wallis test, followed by Dunn’s post-hoc test, was performed. The growth of cells on solid medium was evaluated using dilution spot assays. Bacteria were grown overnight in 5 ml of LB-Km medium. 10-fold serial dilutions of the cultures were spotted as 5-µl drops onto LB Km agar plates supplemented with 0,5 mM IPTG and different concentrations of NaF (0-30 mM NaF). Plates were incubated at 30 °C for 20 hours. Two technical replicate plates were done for each NaF concentration.

### 2.8 Proteomic and transcriptomic analysis

Proteomic and transcriptomic samples were prepared from cultures grown in the same conditions. For proteomic analysis, three and for the transcriptomic analysis, two independent cultures of each *P. putida* strain were sent to be analysed. As the lag-phase is the growth parameter most affected by the fluoride stress (32), to obtain similar effects, all strains were grown in different NaF concentrations that had a similar impact on lag-phase (the lag-phase was 2 times longer than without NaF). For the ΔcrcB strain, 0.2 mM NaF was used; for the ΔcrcBΔ3125 strain, 3.5 mM NaF was used; for the Δ3125 strain, 30 mM NaF was used; and for the WT strain, 33 mM NaF was used. Cells were grown overnight and diluted to fresh media to an OD580 of 0.05. Cultures were grown in biological duplicates until the beginning of the exponential phase, with an OD580 of 0.1, and biomass was harvested and washed twice with 1x M9 buffer (42 mM KH2PO4, 24 mM Na2HPO4, 19 mM NH4Cl, 9 mM NaCl). Cell pellets corresponding to 1 ml of OD580 = 1 were flash-frozen in liquid nitrogen and stored at -80°C until further use.

For proteomics, label-free quantification of the whole cell proteome was performed by LC-MS/MS with LTQ-Orbitrap XL (Thermo Fisher Scientific) coupled to an Agilent 1200 nanoflow LC via nanoelectrospray ion source (Proxeon) in the Proteomics Core Facility, Institute of Technology, University of Tartu, Estonia. The data were analysed using MaxQuant and Perseus software (Max Planck Institute of Biochemistry, Planegg, Germany) (33). Parallel samples were grouped, and groups were compared in pairs: deletion strains against WT strains and parallel with NaF against parallels without NaF. To be included in the analysis, a protein must be detected in all three replicates of one group. Thereafter, missing values were imputed using default settings. Mean protein abundances were compared between the two groups using the independent-sample Student t-test. The Benjamini-Hochberg multiple-testing correction was applied with the false discovery rate set to 0.05. The mass spectrometry proteomics data have been deposited to the ProteomeXchange Consortium via the PRIDE (34) partner repository with the dataset identifier PXD068884.

For transcriptomics, the cultures were prepared the same way as for the proteomic analysis. RNA extraction and purification were performed by the Global Genomic Centre in Hong Kong, China. RNAseq analysis was performed to investigate transcriptomic responses under NaF toxicity conditions. Raw sequencing data were processed using the DESeq2 package (v1.48.1) within R (v4.5.1), applying standard pipelines for normalisation, differential expression analysis, and statistical significance testing. Differentially expressed genes were determined using DESeq2’s default parameters, with p-values adjusted via the Benjamini-Hochberg method to control for false discovery rate. For data visualisation and enhanced customisation of principal component analysis (PCA), MA plots, and volcano plots, the transcriptomics version of the VisomX R package (available at https://github.com/NicWir/VisomX) was employed. The data have been deposited in NCBI’s Gene Expression Omnibus (35) and are accessible through GEO Series accession number GSE309641 (https://www.ncbi.nlm.nih.gov/geo/query/acc.cgi?acc=GSE309641).

## 3. RESULTS

### 3.1 Genomic changes in spontaneous NaF^+^ mutants in the **Δ***crcB* strain

The ability of *P. putida* to tolerate high fluoride concentrations relies on the activity of the CrcB transporter protein as the only known mechanism so far. On LB agar plates, wild-type KT2440 cells can tolerate fluoride concentrations even higher than 45 mM, whereas the strain lacking the CrcB transporter (Δ*crcB* strain) shows growth restrictions already at 0.25 mM of NaF (36). However, we observed the appearance of single colonies when the Δ*crcB* strain was incubated on LB plates containing 5 mM NaF. The spontaneous NaF-tolerant mutant frequency of the Δ*crcB* strain was estimated as specified in Materials and Methods. We observed that the median number of NaF-tolerant mutants that emerged at 5 mM NaF was 5 CFUmL^-1^. Four isolated mutants designated *crcB1*, *crcB4*, *crcB12*, and *crcB18* were sequenced to investigate the mechanisms underlying fluoride tolerance.

Comparative genomic analysis revealed that all four Na-tolerant Δ*crcB* mutants carried large deletions in their genomes, ranging from 18 kb to 340 kb (corresponding to 18-345 genes) as illustrated in Fig. 1. While the deleted regions overlapped, their start and end points varied. The specific deleted areas were as follows: *crcB1* [*PP_2985*]-*PP_5598*; *crcB4* [*PP_2985*]-[*PP_3288*]; *crcB12* [*PP_3119*]-[*PP_3133*]; and *crcB18* [*PP_3059*]-[*PP_3221*], where genes with square brackets are partially deleted. The list of all the deleted genes of the shortest deletion can be found in supplementary table S1.

**Figure 1.**
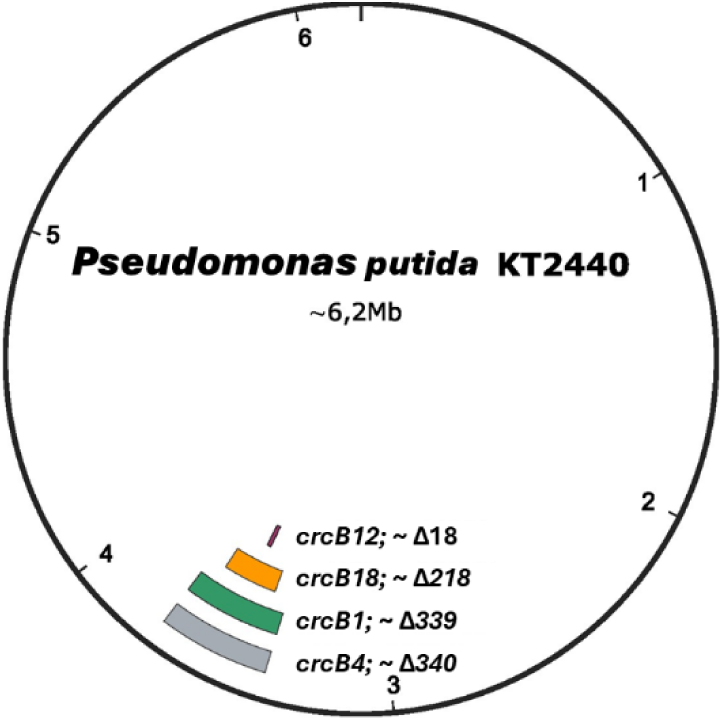
Genomic deletions in isolated spontaneous NaF-tolerant Δ*crcB* mutants *crcB1*, *crcB4*, *crcB12*, and *crcB18*. The different genomic deletions are shown with the isolate number and the deletion size in kilobases.

Given the large deletions, which made it challenging to pinpoint the exact genes responsible for increased fluoride tolerance, we performed transposon mutagenesis using the mini-Tn*5* system to identify specific genes affecting fluoride tolerance in the Δ*crcB* strain.

3.2 Identification of genes affecting fluoride tolerance in *P. putida* **Δ***crcB* strain

To identify the genes affecting fluoride tolerance in *P. putida*, transposon mutagenesis was performed with the Δ*crcB* strain. Approximately 141,000 transposon mutants from three independent transposon mutagenesis experiments were obtained, and the fluoride tolerance of the mutants was analysed on plates containing 5 mM NaF. Out of 141,000 mutants, 48 had developed higher fluoride tolerance. The chromosomal location of mini-Tn*5* was identified in 24 mutants. Mutations were located all over the genome, as shown in Fig. 2 (A), which indicates that there was no significant positional bias in the assay. As seen in Fig. 2 (B, C), from those 24 mutants analysed, 16 contained the transposon insertions into the *PP_3125* gene. We focused on the *PP_3125* gene for further analysis, as all the other insertions targeted the other genes only once. It is worth noting that the *PP_3125* gene was always included in the DNA region that was deleted in the spontaneous NaF-tolerant Δ*crcB* mutants *crcB1*, *crcB4*, *crcB12*, and *crcB18*, shown in Fig. 1.

**Figure 2.**
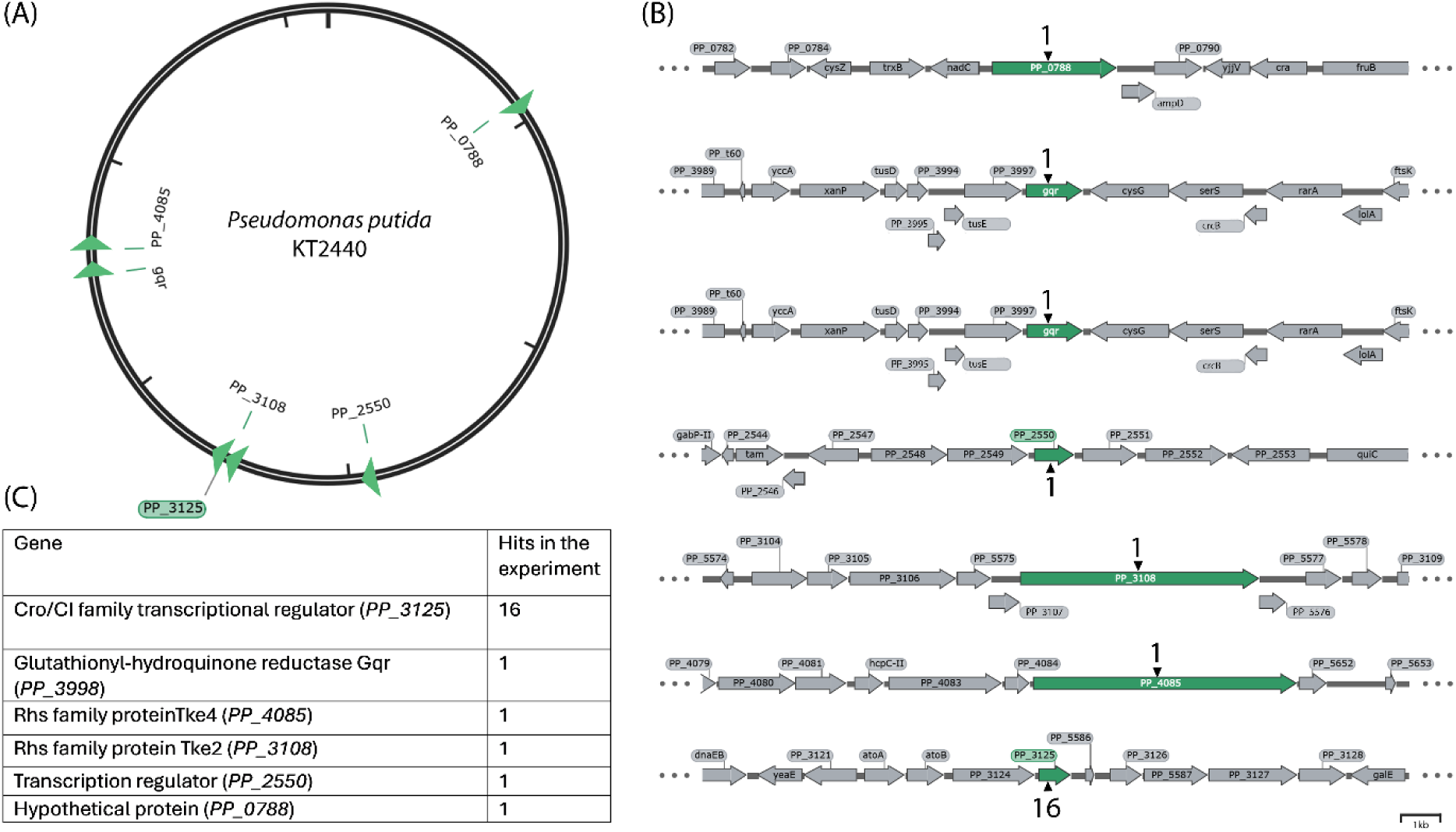
(A) Physical map of the genome of *P. putida* KT2440 illustrating the insertion sites of the transposon. The gene with the most hits, PP_3125, is marked with a green background. (B) Genetic context of transposon insertions and the count of transposon mutants. Genes containing transposon insertions are marked in green, the number of insertions is indicated with black arrowheads, and grey-marked genes represent the genetic context surrounding the genes with transposon insertions. (C) Isolated transposon mutants with gene descriptions and hits per gene.

Our results indicated that inactivation of *PP_3125* encoding Cro/CI type transcriptional regulator increases the fluoride tolerance of the Δ*crcB* strain.

### 3.3 Involvement of the *PP_3125* gene in NaF tolerance of the Δ*crcB* strain

To confirm the effect of PP_3125 in fluoride tolerance, the following strains were constructed: Δ*3125* - *P. putida* KT2440 strain where *PP_3125* gene is deleted; Δ*crcB*Δ*3125* - *P. putida* KT2440 double mutant where both the genes *crcB* and *PP_3125* were deleted; Δ*crcB*Δ*3125*-tac*3125* - double deletion strain with the gene cassette containing then *PP_3125* gene under the control of P*_tac_*promoter in the chromosome at the Tn*7* insertion site. The growth experiments in liquid media and on LB agar plates with different NaF concentrations were carried out. To analyse the growth of constructed strains on solid media, 10-fold serial dilutions of overnight cultures of *P. putida* strains were spotted on LB agar plates with NaF concentrations from 0 to 5 mM NaF and analysed after 20 h of growth. The liquid media growth experiments were carried out for 24 hours under varying NaF concentrations (0, 0.5, and 2.5 mM NaF).

As shown in Fig. 3 and Fig. 4, in the absence of NaF, all strains exhibited similar growth patterns. However, in the presence of 0.5 mM NaF, the Δ*crcB* strain showed reduced growth on LB agar plates, whereas the Δ*crcB*Δ*3125* strain did not show any growth restrictions (Fig. 3). Also, in liquid media, at 0.5 mM NaF, the Δ*crcB* strain showed a longer *lag-*phase than Δ*crcB*Δ*3125* (10.732 h vs 1.56 h, adjusted p value = 0.0293). With the higher fluoride concentrations, the differences were even bigger. The difference in maximum growth rate was not relevant, which shows that exposure of these strains to fluoride affects the length of the *lag*-phase but not the growth rate.

**Figure 3.**
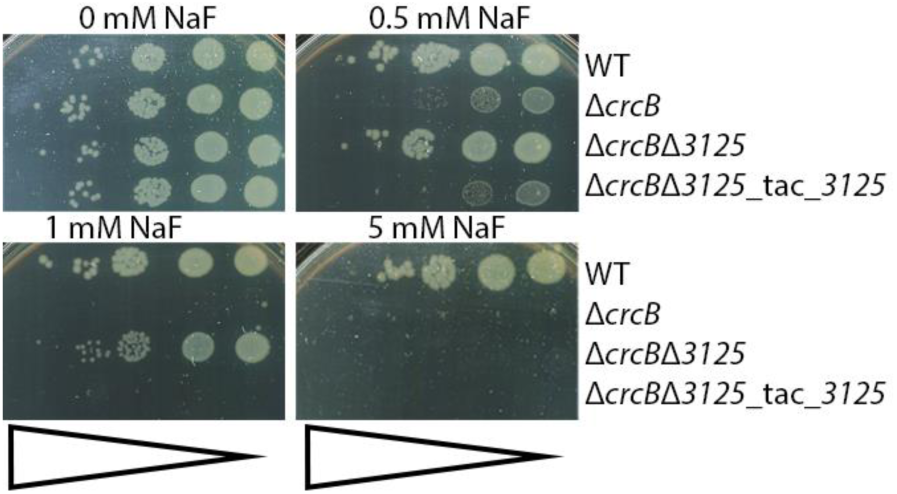
Dilution spot assay of *P. putida* WT, Δ*crcB*, Δ*3125*, Δ*crcB*Δ*3125,* and Δ*crcB*Δ*3125*-tac*3125* strains on LB plates supplemented with 0-5 mM NaF. Growth on different fluoride concentrations was examined by spotting 10-fold serial dilutions (indicated by bars) of overnight cultures of the strains onto plates with different NaF concentrations. The plates were incubated at 30°C for 20 h.

**Figure 4.**
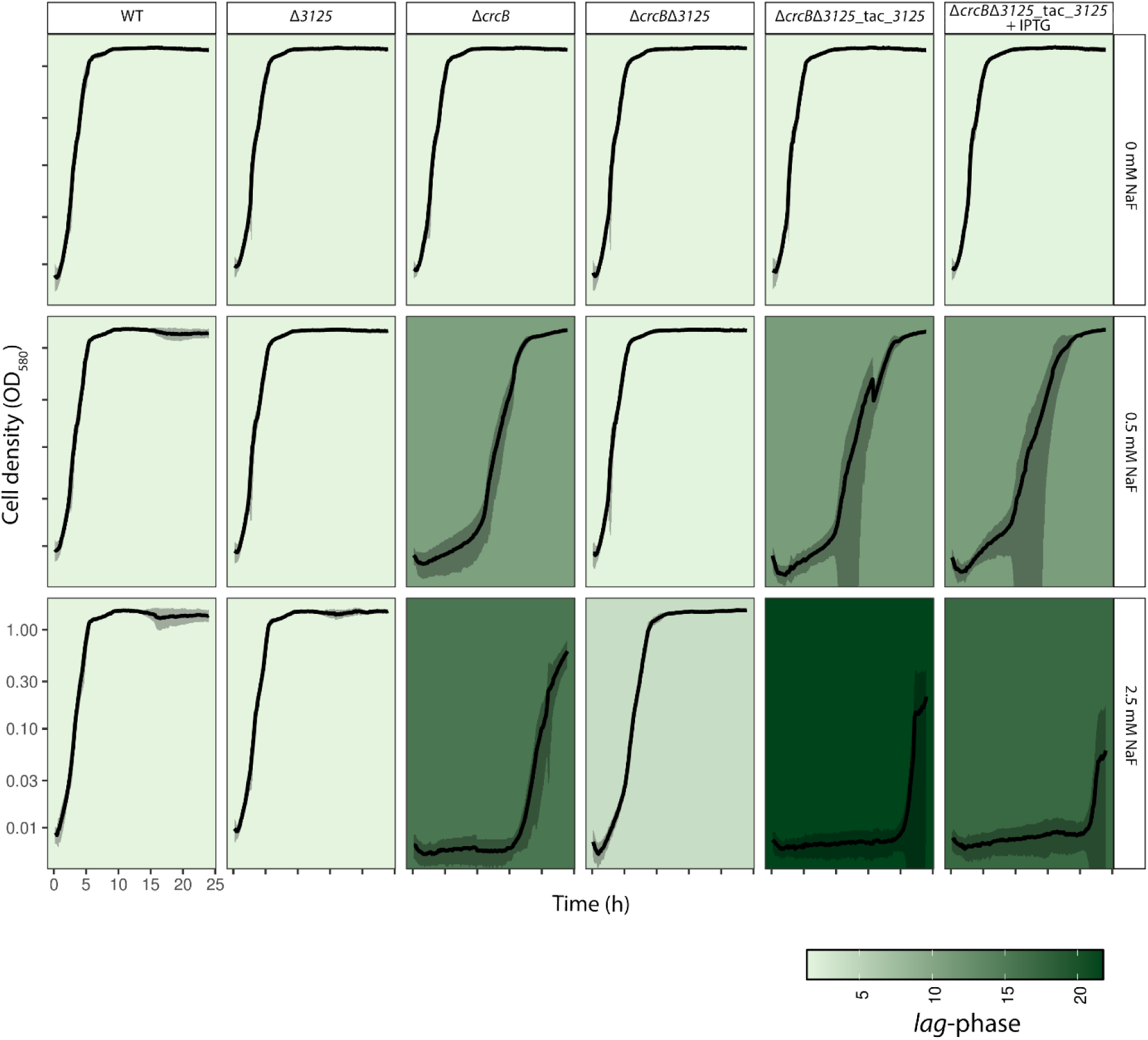
Growth curves of *P. putida* WT, Δ*crcB*, Δ*3125*, Δ*crcB*Δ*3125,* and Δ*crcB*Δ*3125*-tac*3125* strains growing on different NaF concentrations. 0.01 mM IPTG was used to induce the *tac*-promoter. Averages of three different biological replicas with three different technical parallels are presented with standard deviation. The background colours show the length of the lag-phase; the darker the background, the longer the lag-phase.

Due to leaky expression from the P*_tac_* promoter, the Δ*crcB*Δ*3125*-tac*3125* strain showed similar growth to the Δ*crcB* strain in NaF-containing media, even without IPTG induction (Fig. 3, Fig. 4, Table 3). The high variability in growth curves of fluoride-sensitive strains in liquid media at the end of the experiment could be attributed to the emergence of spontaneous mutants capable of tolerating elevated fluoride concentrations.

**Table 3.**
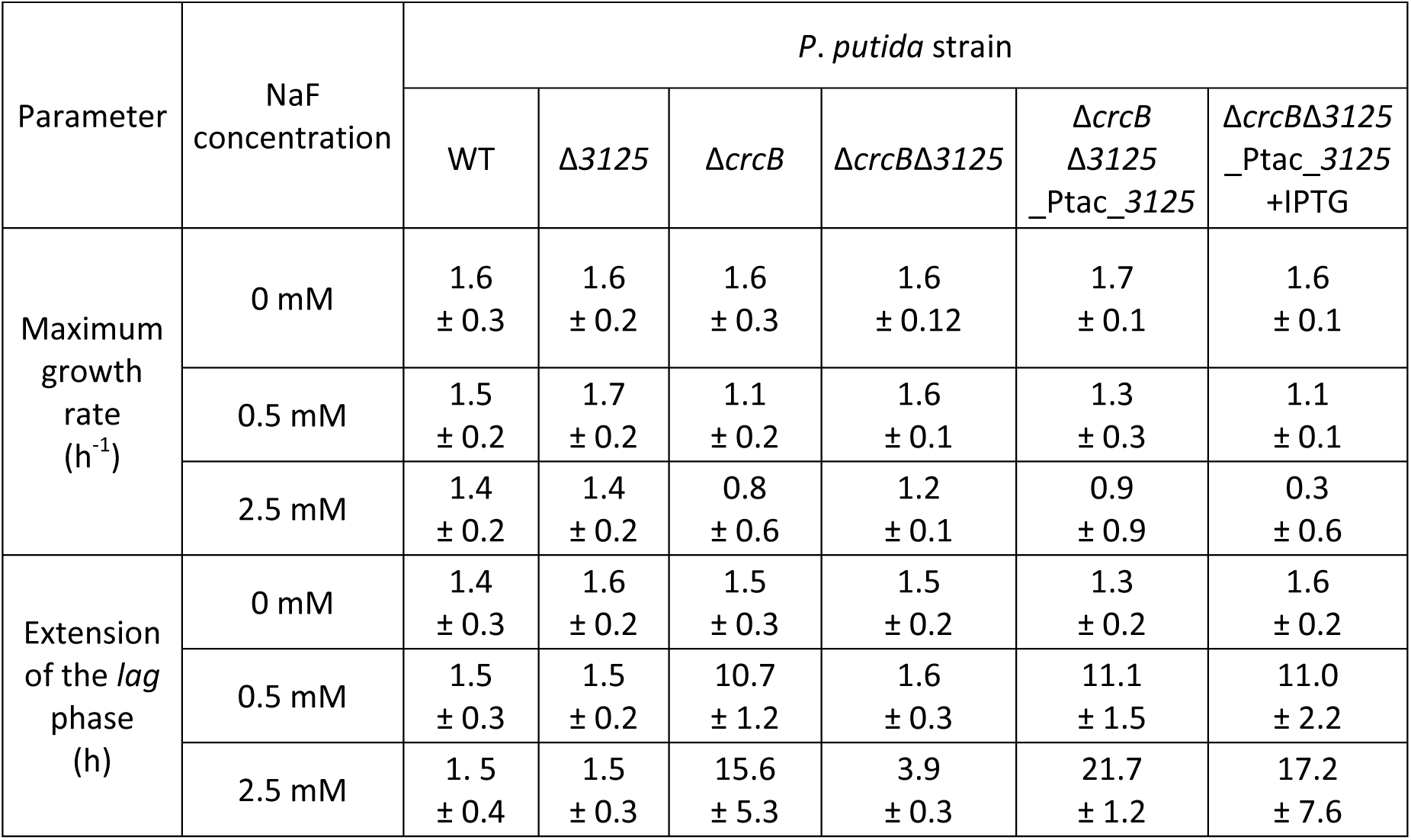
Maximum growth rate and *lag-*phase of *P. putida* WT, Δ*crcB*, Δ*3125*, Δ*crcB*Δ*3125,* and Δ*crcB*Δ*3125*-tac*3125* strains growing on different NaF concentrations.

### 3.4 Spontaneous NaF-tolerant Δ*crcB* mutants have similar growth characteristics to the Δ*crcB*Δ*3125* strain

Since *PP_3125* is located within the deleted regions of the spontaneously occurring NaF-tolerant Δ*crcB* mutants, we investigated whether the deletion of *PP_3125* alone had a similar effect on NaF tolerance as that observed in the spontaneous mutants.

As shown in Fig. 5 and Table 4, the spontaneous mutants exhibited growth comparable to the wild-type (WT) strain in the absence of NaF. However, when NaF was added, their growth patterns closely resembled the growth of the Δ*crcB*Δ*3125* strain. When the *lag-*phase of the Δ*crcB* strain with 2.5 mM NaF was 18.925 h long, the *lag-*phase of the spontaneous mutants and the Δ*crcB*Δ*3125* strain was 7 times shorter, but the adjusted p-values weren’t significant. This might be due to a very high variability in the growth pattern of the Δ*crcB.* But the results still indicate that the NaF tolerance observed in the spontaneous mutants is most probably due to the absence of *PP_3125*.

**Figure 5.**
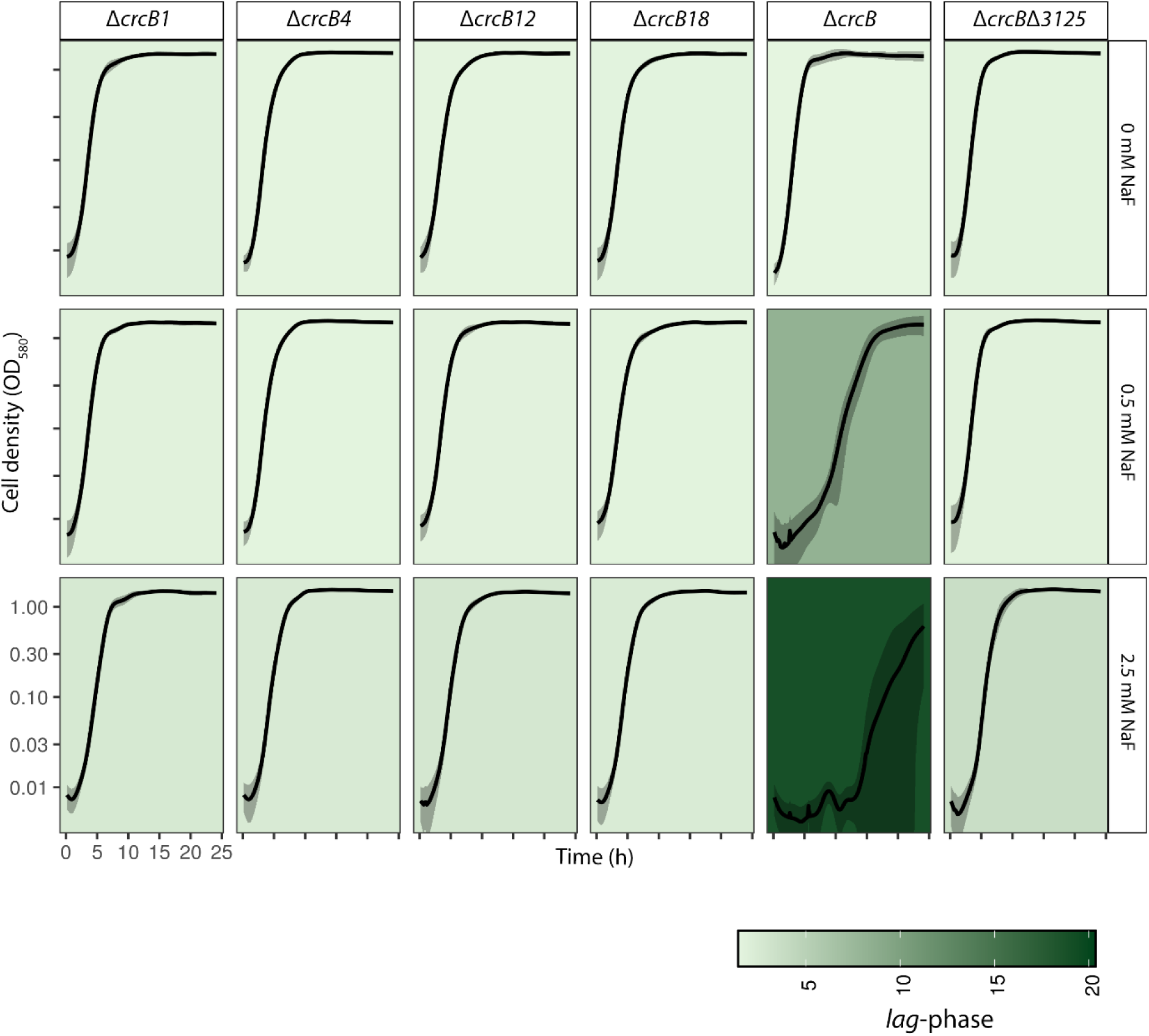
Growth curves of *P. putida* Δ*crcB*, Δ*crcB*Δ*3125*, and spontaneous NaF+ ΔcrcB mutants crcB1, 12, 18, and 4 growing on different NaF concentrations. Averages of three different biological replicas with three different technical parallels are presented with standard deviation. The background colours show the length of the *lag-*phase; the darker the background, the longer the *lag-*phase.

**Table 4.** Maximum growth rate and *lag-*phase of *P. putida* Δ*crcB*, Δ*crcB*Δ*3125*, and spontaneous NaF+ ΔcrcB mutants crcB1, 12, 18, and 4 growing on different NaF concentrations.

| Parameter | NaF concentration | *P. putida* strain |  |  |  |  |  |
| --- | --- | --- | --- | --- | --- | --- | --- |
| | | Δ*crcB*1 | Δ*crcB*4 | Δ*crcB*12 | Δ*crcB*18 | Δ*crcB* | Δ*crcB*Δ3125 |
| Maximum growth rate (h<sup>-1</sup>) | 0 mM NaF | 1.5<br>± 0.2 | 1.5<br>± 0.1 | 1.4<br>± 0.2 | 1.5<br>± 0.2 | 1.6<br>± 0.1 | 1.6<br>± 0.2 |
|  | 0.5 mM NaF | 1.5<br>± 0.2 | 1.5<br>± 0.2 | 1.4<br>± 0.1 | 1.4<br>± 0.1 | 1.0<br>± 0.2 | 1.6<br>± 0.2 |
|  | 2.5 mM NaF | 1.3<br>± 0.2 | 1.3<br>± 0.1 | 1.4<br>± 0.2 | 1.3± 0.2 | 1.0<br>± 0.5 | 1.5<br>± 0.2 |
| Extension of *lag*-phase (h) | 0 mM NaF | 1.8<br>± 0.1 | 1.5<br>± 0.1 | 1.5<br>± 0.2 | 1.7<br>± 0.3 | 1.5<br>± 0.1 | 1.7<br>± 0.1 |
|  | 0.5 mM NaF | 1.6<br>± 0.2 | 1.6<br>± 0.1 | 1.7<br>± 0.1 | 1.7<br>± 0.3 | 8.2<br>± 1.1 | 1.8<br>± 0.2 |
|  | 2.5 mM NaF | 2.7<br>± 0.4 | 2.5<br>± 0.1 | 2.8<br>± 0.3 | 2.4<br>± 0.2 | 18.9<br>± 3.4 | 3.7<br>± 0.4 |

### 3.5 Identification of genes regulated by PP_3125

A proteomic analysis was conducted to identify the genes regulated by PP_3125 that might contribute to fluoride tolerance. All strains were grown under different NaF concentrations, resulting in similar growth effects, specifically a doubling of the *lag-*phase compared to the growth without NaF. For the Δ*crcB* strain, 0.2 mM NaF, for the Δ*crcB*Δ*3125* strain, 3.5 mM NaF, for the Δ*3125* strain, 30 mM NaF, and for the WT strain, 33 mM NaF concentrations were used.

As shown in Fig. 6A, in the presence of NaF, only three proteins exhibited significantly differential expression when comparing the *PP_3125* deletion strain to the WT strain in the presence of fluoride. The overproduced proteins were PP_2036 (4-hydroxy-tetrahydrodipicolinate synthase), PP_2037 (aldolase), and BenE-I (benzoate transport protein). As these same protein genes were also overexpressed in the Δ*crcB*Δ*3125* strain compared to both the WT (Supplementary Fig. S1) and the ΔcrcB strain in the presence of NaF (Supplementary Fig. S2), this suggests that NaF may regulate *PP_3125* expression.

**Figure 6.**
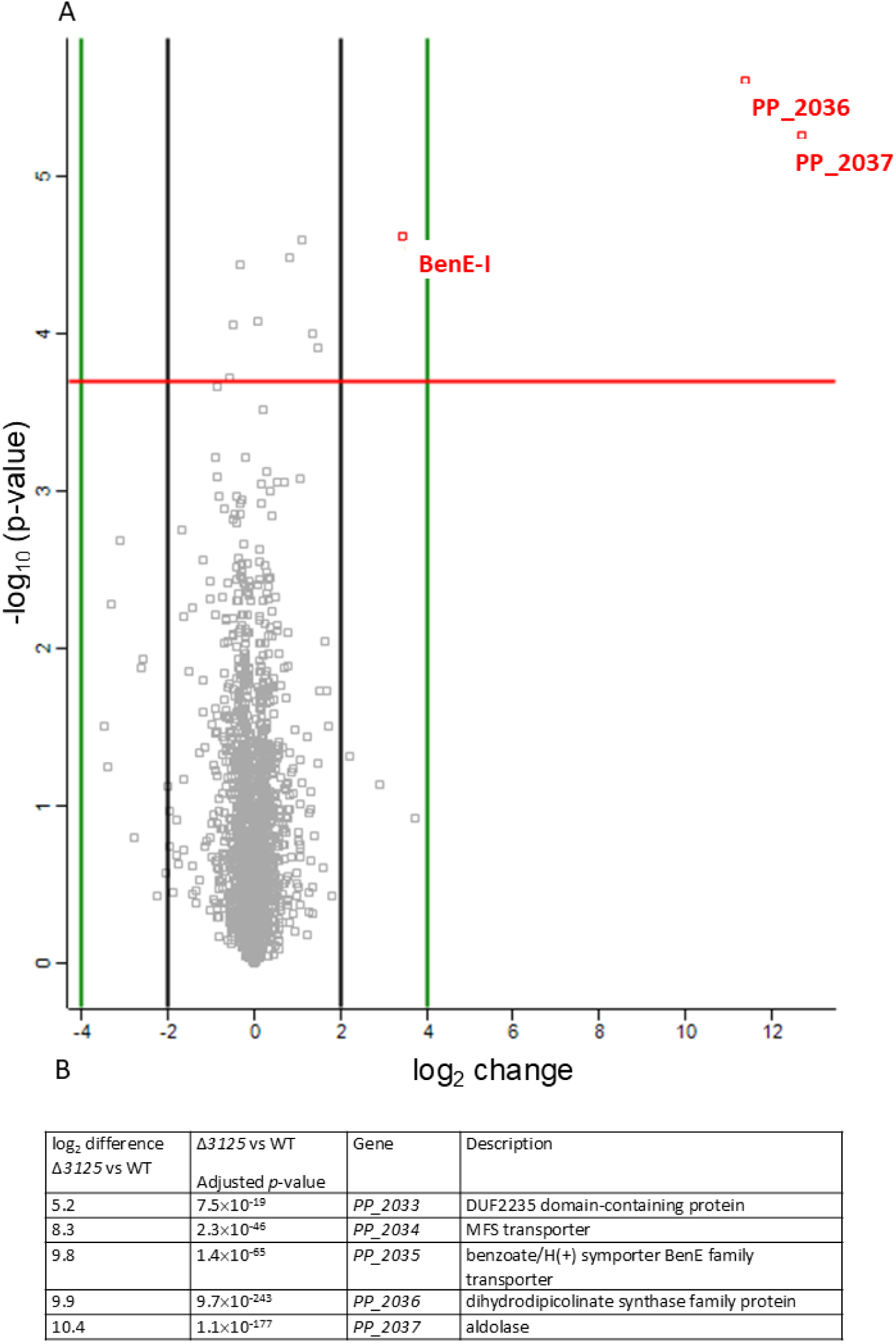
Overview of *P. putida* WT and Δ*3125* strains’ full proteome and transcriptome comparison in the presence of NaF. A. In a volcano plot, every dot represents a protein. The horizontal line indicates the statistical significance threshold after Benjamin–Hochberg multiple testing correction [false discovery rate (FDR) = 0.05]. The vertical lines indicate a twofold difference between the compared proteomes. Statistically significant at least two-fold increase in protein level are presented as red dots with the gene name. B. Transcriptome analysis of *P. putida* genes in the absence of PP_3125.

To further explore which other genes could also be regulated by PP_3125, the transcriptome of the WT, Δ*crcB*, Δ*3125,* and Δ*crcB*Δ*3125* strains was prepared under the same conditions as those performed in proteome studies.

Transcriptome analysis revealed the distinct effects of fluoride, *crcB* deletion, and *PP_3125* deletion (Fig. 6B). When PP_3125 proficient strains were compared to the strain lacking the regulator PP_3125 in the presence of fluoride in the growth medium, we were able to confirm only the differential expression of 5 genes. Other differently expressed genes resulted from the fluoride stress (Supplementary Table S2). Five genes overexpressed in the absence of *PP_3125* are in the same genomic region, adjacent to each other. This indicated that these genes could be co-regulated. Importantly, among the five identified genes, *PP_2036*, *PP_2037*, and *benE-I* (*PP_2035*) were also overexpressed in the proteomic analysis. The other two genes, *PP_2034* (a hypothetical transporter) and *PP_2033* (of unknown function), were clustered in the same region. The results of the transcriptome analysis can be found in the supplementary table S3.

### 3.7 PP_3125-regulated genes affecting fluoride tolerance

To determine which of the five genes identified in the transcriptomic study could affect fluoride tolerance, a deletion strain lacking all five genes (PP_2033-PP_2037) was first constructed. As two out of 5 genes in the operon were transporters (*PP_2034* and *benE-I*), we suspected that one of the transporters could also transport fluoride in addition to its primary substrate. We first constructed plasmid-based overexpression constructs of these two genes, where these are under the control of a P*_tac_* promoter.

Due to leaky expression from the P*_tac_* promoter, the effect of the transporters was observed even in the absence of IPTG. As shown in Fig. 7 and Table 5, the Δ*crcB*Δ5 strain lacking all five genes identified in the transcriptomics study (*PP_2033-PP_2037)* could not grow in the presence of 2.5 mM NaF, even after 24 hours. Introducing the *PP_2034* transporter to the Δ*crcB*Δ5 strain resulted in minimal growth toward the end of the experiment; however, this growth was minimal. In contrast, introducing the BenE-I transporter significantly improved the fluoride tolerance of the Δ*crcB*Δ5 strain, allowing growth at 2.5 mM NaF. Although expressing *benE-I* did not restore the fluoride tolerance to the wild-type levels, the tolerance was notably higher than that of the Δ*crcB* strain.

**Figure 7.**
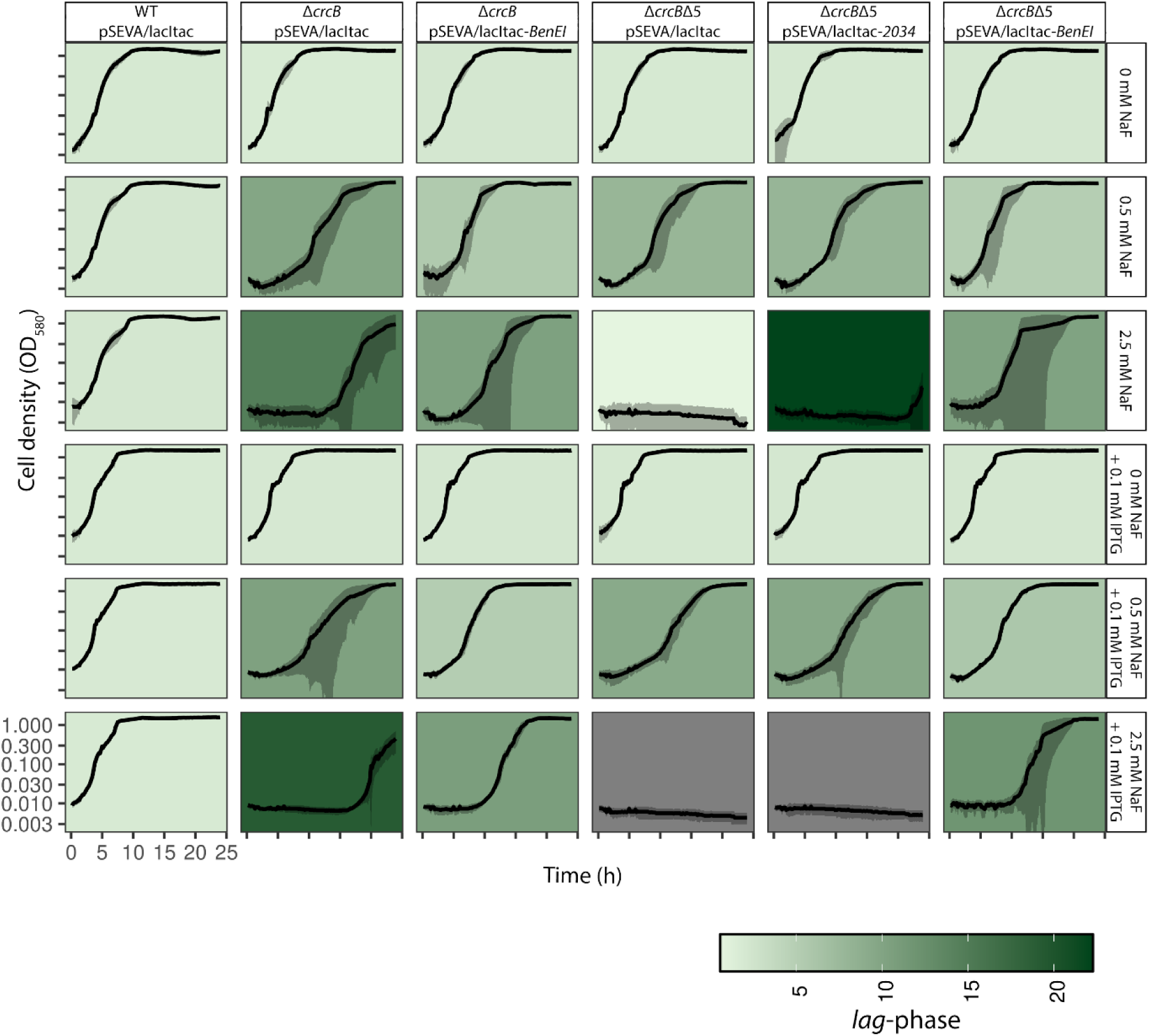
Growth curve of *P. putida* WT, Δ*crcB,* and Δ*crcB*Δ*5* with empty pSEVA/lacItac plasmid or with *PP_2034* or *benE-I* in a plasmid under the control of P*_tac_* growing on different NaF concentrations and with or without 0.1 mM IPTG. Averages of three different biological replicas with three different technical parallels are presented with standard deviation. The background colours show the length of *lag-*phase, the darker the background, the longer the *lag-*phase. 0.1 mM of IPTG is added as an inductor.

**Table 5.** Maximum growth rate and *lag-*phase of *P. putida* WT, Δ*crcB,* and Δ*crcB*Δ*5* with empty pSEVA/lacItac plasmid or with *PP_2034* or *benE-I* in a plasmid under the control of P*_tac_* growing on different NaF concentrations and with or without 0.1 mM IPTG.

| Parameter | NaF<br>concentration | *P. putida* strain |  |  |  |  |  |
| --- | --- | --- | --- | --- | --- | --- | --- |
|  |  | WT<br>pSEVA/lacIac | *ΔcrcB*<br>pSEVA/lacIac | *ΔcrcB*<br>pSEVA/lacIac<br>-*BenEI* | *ΔcrcBΔ5*<br>pSEVA/lacIac | *ΔcrcBΔ5*<br>pSEVA/lacIac<br>-2034 | *ΔcrcBΔ5*<br>pSEVA/lacIac<br>-*BenEI* |
| Maximum growth rate (h<sup>-1</sup>) | 0 mM NaF | 1.2 ± 0.1 | 1.2 ± 0.1 | 1.1 ± 0.1 | 1.2 ± 0.1 | 1.2 ± 0.2 | 1.2 ± 0.1 |
|  | 0.5 mM NaF | 1.1 ± 0.2 | 1.0 ± 0.1 | 1.4 ± 0.1 | 1.2 ± 0.2 | 1.2 ± 0.1 | 1.3 ± 0.1 |
|  | 2.5 mM NaF | 1.1 ± 0.1 | 0.9 ± 0.7 | 1.4 ± 0.3 | 0.1 ± 0.2 | 0.4 ± 0.5 | 1.1 ± 0.1 |
|  | 0 mM NaF + 0.1 mM IPTG | 1.2 ± 0.1 | 1.4 ± 0.1 | 1.4 ± 0.0 | 1.3 ± 0.2 | 1.4 ± 0.1 | 1.4 ± 0.1 |
|  | 0.5 mM NaF + 0.1 mM IPTG | 1.2 ± 0.0 | 1.0 ± 0.2 | 1.1 ± 0.0 | 0.9 ± 0.1 | 1.0 ± 0.2 | 1.1 ± 0.1 |
|  | 2.5 mM NaF + 0.1 mM IPTG | 1.2 ± 0.1 | 1.2 ± 0.4 | 1.2 ± 0.1 | 0 | 0 | 1.1 ± 0.1 |
| Extension of lag-phase (h) | 0 mM NaF | 2.0 ± 0.2 | 1.8 ± 0.2 | 2.2 ± 0.2 | 2.0 ± 0.2 | 2.1 ± 0.7 | 2.2 ± 0.3 |
|  | 0.5 mM NaF | 2.1 ± 0.1 | 9.9 ± 1.6 | 5.6 ± 0.7 | 7.9 ± 1.1 | 8.0 ± 0.8 | 5.1 ± 1.2 |
|  | 2.5 mM NaF | 2.2 ± 0.5 | 14.7 ± 1.0 | 10.4 ± 2.6 | 0.6 ± 0.7 | 22.2 ± 0.5 | 10.2 ± 3.8 |
|  | 0 mM NaF + 0.1 mM IPTG | 1.9 ± 0.1 | 2.0 ± 0.2 | 2.2 ± 0.1 | 2.1 ± 0.2 | 2.2 ± 0.1 | 2.2 ± 0.1 |
|  | 0.5 mM NaF + 0.1 mM IPTG | 1.9 ± 0.1 | 10.4 ± 3.6 | 5.2 ± 0.5 | 9.4 ± 1.3 | 9.6 ± 1.6 | 5.9 ± 0.1 |
|  | 2.5 mM NaF + 0.1 mM IPTG | 1.9 ± 0.1 | 19.2 ± 1.4 | 10.7 ± 0.2 | 0 | 0 | 11.9 ± 1.4 |

Additionally, when the BenE-I transporter-encoding gene under the P*_tac_* promoter control was introduced into the Δ*crcB* strain, a shorter lag phase was observed compared to the Δ*crcB* strain without the BenE-I transporter.

The addition of IPTG into growth medium of bacteria further amplified the impact of the five-gene deletion, leading to even more pronounced growth inhibition by NaF. However, we observed that IPTG itself affected bacterial growth, as the Δ*crcB* strain with an empty plasmid exhibited a more extended lag phase in the presence of IPTG compared to growth without IPTG supplementation (14.746 h without IPTG vs 19.207 h with IPTG; however, no statistically significant difference was observed).

To compare the growth of *P. putida* KT2440 wild-type strain, Δ*crcB* strain, and its derivatives when either the *PP_3125* regulator gene or five genes identified in a transcriptomics study (*PP_2033-PP_2037)* were removed additionally, and the strains with the plasmid containing a *benE-I* overexpression construct, a growth assay on LB agar plates was carried out. All the compared strains contained either an empty pSEVA/lacItac plasmid or a pSEVA/lacItac-*benE-I* plasmid. The results shown in Fig. 8 demonstrated that all strains grew equally well on LB media without NaF. Strains lacking the CrcB transporter did not grow on 5 mM NaF, whereas the WT strain could tolerate over 30 mM NaF. Removing transcription regulator PP_3125 increased fluoride tolerance of the Δ*crcB* strain, as the strain missing both CrcB and PP_3125 was able to grow on 5 mM NaF as well as the wild-type strain. As indicated by the growth in liquid media (Fig. 7), PP_3125 could be the negative regulator of BenE-I. When this regulator is absent, the expression of the *benE-I* transporter is higher, and bacteria can tolerate higher concentrations of NaF. Moreover, we observed that when the BenE-I transporter was artificially overproduced under the control of the *P_tac_* promoter in the Δ*crcB* strain or in the Δ*crcB*Δ5 strain, the NaF tolerance increased further, allowing bacterial growth even at 20 mM NaF (Fig. 8).

**Figure 8.**
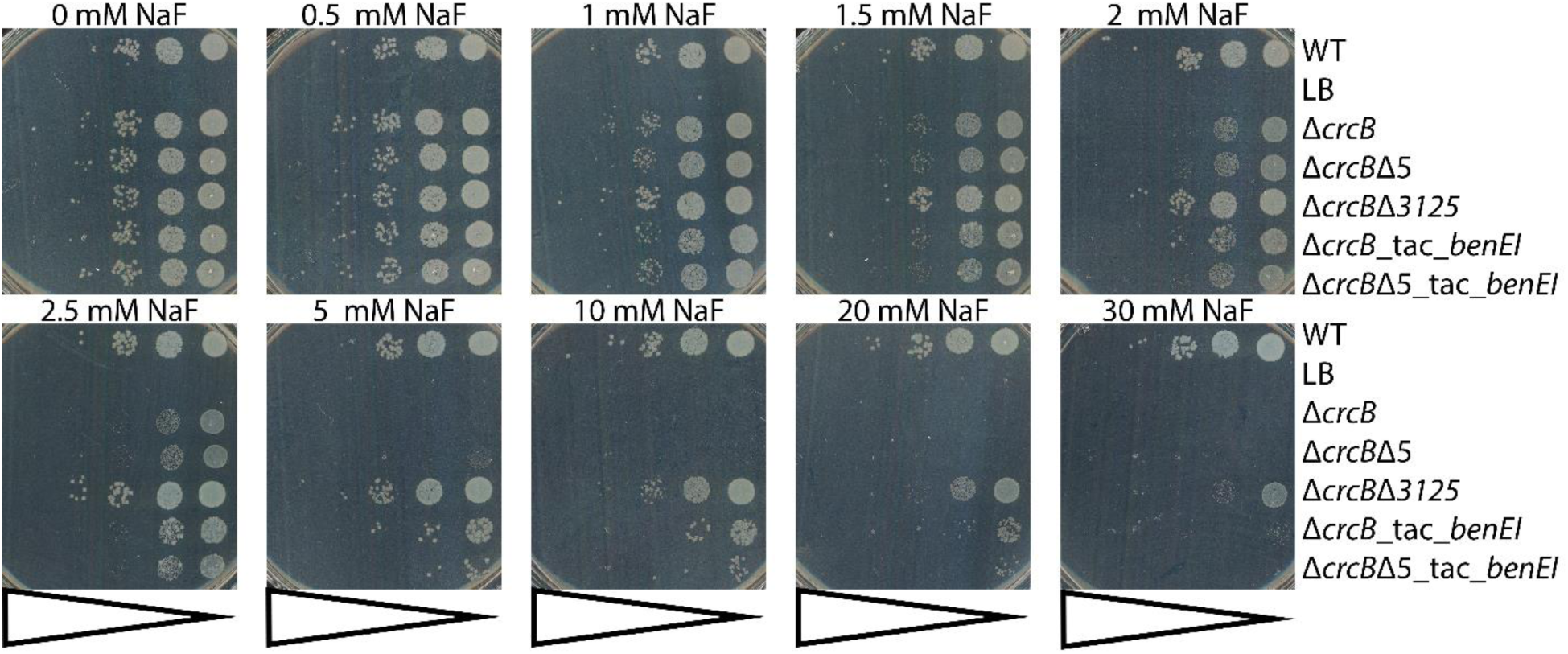
Dilution spot assay of *P. putida* WT, Δ*crcB*Δ5, Δ*crcB*Δ*3125,* Δ*crcB_*tac_*benEI,* and Δ*crcB*Δ5_tac_*benEI* on LB-Km plates supplemented with 0-30 mM NaF and 0.1 mM IPTG as P_tac_-promoter inducer. Growth on different fluoride concentrations was examined by spotting 10-fold serial dilutions (indicated by bars) of overnight cultures of the strains onto plates with different NaF concentrations. The plates were incubated at 30°C for 20 h.

Overall, these results demonstrated that PP_3125 functions as a negative regulator of a gene influencing fluoride tolerance in *P. putida*. BenE-I, a benzoate transporter regulated by PP_3125, enhances fluoride tolerance when overexpressed. Naturally occurring NaF-tolerant Δ*crcB* mutants appeared to gain their tolerance through deletion of a genomic region containing *PP_3125*, as this gene was absent in all sequenced spontaneous mutants, and the fluoride tolerance levels of spontaneous mutants and the Δ*crcB*Δ*3125* deletion strain were comparable.

## 4. Discussion

The findings of this study significantly advance our understanding of fluoride tolerance mechanisms in *P. putida*. While previous studies highlighted the role of the CrcB transporter (also known as Fluc) in exporting intracellular fluoride ions (6). This work uncovers an additional, CrcB-independent pathway involving the transcriptional regulator PP_3125. Deletion of *PP_3125* in the Δ*crcB* background led to a marked increase in fluoride tolerance (Fig. 4), mirroring the phenotype observed in spontaneous fluoride-tolerant mutants with large genomic deletions encompassing this gene (Fig. 5).

There are two distinct classes of fluoride transporters: ClCF and CrcB (Fluc) transporters (4,5,37). ClCF transporters function as proton-fluoride antiporters, facilitating active fluoride extrusion from the cell (37). In contrast, CrcB transporters operate as passive fluoride-specific ion channels, requiring Na⁺ ions for transport activity (3,4,38–41). While ClCF transporters can transport both chloride and fluoride ions or only fluoride, CrcB transporters are highly selective for fluoride (5,42). These mechanistic distinctions underscore the importance of identifying alternative fluoride tolerance strategies when CrcB is deleted or nonfunctional.

Using transposon mutagenesis, we identified *PP_3125* as the most frequently disrupted gene in fluoride-tolerant Δ*crcB* mutants, indicating its central role as a negative regulator of the fluoride stress response. Proteomic and transcriptomic analyses further revealed that PP_3125 negatively regulates a cluster of five adjacent genes (*PP_2033-PP_2037*), including transporters and metabolic enzymes. Notably, the BenE-I transporter, a member of the benzoate/H⁺ symporter family, emerged as a key factor in promoting fluoride tolerance when overexpressed. Restoration of fluoride tolerance in Δ*crcB* strains through overexpression of BenE-I underscores its potential function in either fluoride efflux or maintaining ion homeostasis under fluoride-induced stress (Fig. 7, 8), as the lag-phase of the BenE-I complementation strain is shorter than the deletion strain. The lag-phase is the growth parameter most strongly affected by the fluoride stress (32).

*P. putida* is known for its capacity to metabolize a wide range of aromatic compounds, including benzoate. Key transporters involved in benzoate transport are BenK, BenE, and BenF. BenK and BenE function as aromatic acid-H⁺ symporters, while BenF operates as a benzoate-specific porin and efflux pump (43,44). The functional redundancy and substrate specificity of these transporters reflect the organism’s metabolic adaptability. Given that BenE-I is structurally related to BenE, it is plausible that it shares mechanistic features that allow it to contribute to fluoride tolerance, either directly via efflux or indirectly by modifying membrane potential or ion gradients. The clear mechanism by which the aromatics transporter can also function as a fluoride transporter is still unclear. One of the hypotheses is that BenE-I has an additional N-terminal part, which might play a role in fluoride export. What does the operon *PP_2033*-*PP_2037*, where BenE-I belongs, is unclear, as the aldolase and 4-hydroxy-tetrahydrodipicolinate synthase belong to different metabolic pathways and different steps, and all three other genes have no pathways associated with them. Based on the KEEG database, aldolase PP_2037 is predicted to belong to 6 different pathways: monobactam biosynthesis, lysine biosynthesis, metabolic pathways, biosynthesis of secondary metabolites, microbial metabolism in diverse environments, and biosynthesis of amino acids (45). 4-hydroxy-tetrahydrodipicolinate synthase is predicted to belong to 4 different pathways: ppu00040 Pentose and glucuronate interconversions, fructose and mannose metabolism, metabolic pathways, and microbial metabolism in diverse environments (46).

The lack of clear functional linkage among genes in the PP_2033-PP_2037 operon raises the possibility that their roles in fluoride tolerance may not be confined to canonical metabolic pathways. Instead, they could participate in alternative processes through functional versatility. One intriguing explanation is the phenomenon of protein moonlighting, where a single protein carries out more than one unrelated role. Such multifunctionality has been increasingly recognized in bacteria, where metabolic enzymes, structural proteins, and transporters adopt additional roles beyond their canonical functions (47,48). For example, enolase and glyceraldehyde-3-phosphate dehydrogenase are well-known metabolic enzymes that also act as RNA-binding proteins or surface adhesins in diverse bacterial species (49,50). In the context of fluoride tolerance, the possibility that BenE-I exhibit moonlighting activities cannot be excluded. Given the observed regulatory complexity and the lack of clear pathway associations for most genes in this operon, it is plausible that their encoded proteins may perform secondary functions relevant to ion homeostasis, membrane remodelling, or stress adaptation. This perspective opens the possibility that fluoride tolerance in *P. putida* is not mediated solely through classical efflux but may also involve multifunctional proteins co-opted under selective pressure.

Beyond fluoride-specific responses, *P. putida* encodes a range of transporters with broad substrate specificities that contribute to its environmental resilience. These include heavy metal efflux pumps such as CadA1, CadA2 (P-type ATPases), and CzcCBA1/2 chemiosmotic transporters for Zn²⁺, Cd²⁺, Pb²⁺, Ni²⁺, and Co²⁺ (51), as well as nutrient uptake systems like the AatJMOP complex for glutamate and aspartate (52,53). Efflux systems from the HAE-1 and RND superfamilies, including MexAB-OprM, confer multidrug resistance and can also export dyes and salts (54,55). The outer membrane protein TodX facilitates hydrocarbon transport and exemplifies how *P. putida* adapts to a variety of hydrophobic substrates (35). Such versatility supports the hypothesis that alternate transporters like BenE-I can be co-opted for fluoride tolerance under selective pressure.

The regulatory landscape of PP_3125 may extend beyond what was captured in our proteomic and transcriptomic snapshots, which were limited to early growth phases. It is likely that PP_3125 influences additional gene sets during later growth stages, potentially coordinating broader adaptive responses to prolonged fluoride exposure.

Interestingly, spontaneous fluoride-tolerant mutants showed large genomic deletions encompassing *PP_3125* and up to 344 other genes. These deletions varied in size, from 18 to 345 genes, and included ECF family sigma factors, DNA polymerases (e.g., *dnaEB, imuB*), transporters, regulators, hydrolases, and multiple catabolic pathways such as benzoate and phenylacetate degradation operons. Despite these extensive deletions, the mutants exhibited no significant growth defects under standard laboratory conditions and showed improved growth under fluoride stress (Fig. 5), suggesting that the deleted regions may include nonessential genes under those conditions or that the deletions conferred adaptive benefits.

Multiple mechanisms may underlie these large-scale deletions. Genome plasticity in *P. putida* is supported by the presence of mobile genetic elements such as transposons and prophages. For instance, the *bph-sal* element in *P. putida* KF715 is known to be both deletable and transferable, reflecting high genomic fluidity (56). Moreover, *P. putida* KT2440 contains prophages capable of spontaneous excision, which may promote deletion events even in the absence of phage infectivity (57). It is possible that environmental stresses like fluoride exposure can cause DNA damage, and imprecise repair via the non-homologous end joining (NHEJ) pathway (58) may result in random deletions, but as the deletions are 18-300 kb, there needs to be other mechanisms involved too. The diversity in the start and end points of observed deletions (Fig. 1) further suggests the absence of a single deletion mechanism and highlights the stochastic nature of stress-induced genomic rearrangements.

Taken together, our data support a model in which PP_3125 functions as a central negative regulator of fluoride tolerance, suppressing expression of transporter genes like *benE-I* that mitigate fluoride toxicity. Deletion of *PP_3125*, whether through spontaneous genomic rearrangement or targeted mutagenesis, leads to upregulation of these protective pathways. The fluoride-responsive expression pattern of PP_3125 hints at a yet-to-be-characterized regulatory mechanism that senses and responds to fluoride stress.

In conclusion, this study identifies PP_3125 as a key node in the regulation of fluoride tolerance in *P. putida* and highlights BenE-I as a promising target for engineering enhanced fluoride tolerance. The evolutionary emergence of large deletions encompassing PP_3125 under stress conditions reflects the organism’s remarkable genomic plasticity. Future studies should elucidate the precise function of BenE-I in fluoride tolerance, the broader regulon of PP_3125 across growth phases, and the molecular triggers underlying stress-induced genomic deletions.

## Acknowledgements

This study was supported by the Estonian Research Council (Eesti Teadusagentuur) grant PRG707 to MK, by the European Union’s Horizon 2020 Research and Innovation Programme under grant agreement No. 814418 (SinFonia) and No. 101082049 (TOLERATE), by the The Novo Nordisk Foundation through grants NNF20CC0035580, LiFe (NNF18OC0034818), TARGET (NNF21OC0067996), FM·Pseudomonas (NNF24OC0091501), and NovoF (NNF23OC0083631). The authors gratefully acknowledge Kristo Ets and Jaan Grosberg for their helpful discussions and encouragement throughout the course of this work.

## Data availability

Further information supporting the findings of this study is available upon request from the corresponding author. All the resulting raw sequences from whole genome sequencing are available in GenBank under BioProject ID PRJNA1330567. The mass spectrometry proteomics data have been deposited to the ProteomeXchange Consortium via the PRIDE (34) partner repository with the dataset identifier PXD068884. The transcriptomics data have been deposited in NCBI’s Gene Expression Omnibus (35) and are accessible through GEO Series accession number GSE309641 (https://www.ncbi.nlm.nih.gov/geo/query/acc.cgi?acc=GSE309641).

## Author contributions

Conceived and designed the experiments: L.E, H. I., M.K.

Performed the experiments: L.E., H.I.

Analysed the data: L.E., H.I., T.I., O.P., P.I.N., M.K.

Wrote the initial draft L.E.

Manuscript finalization: L.E., P.I.N., M.K.

